# Ethyl-Iophenoxic acid as a serum marker for oral baiting of carnivorous marsupials

**DOI:** 10.1101/2021.12.14.472710

**Authors:** Ruth Pye, David Nichols, Amy T. Gilbert, Andrew S. Flies

**Affiliations:** Menzies Institute for Medical Research, University of Tasmania, 17 Liverpool St, Hobart, Tasmania 7000, Australia; Central Science Laboratory, College of Sciences and Engineering, University of Tasmania, Sandy Bay, Tasmania 7005, Australia; US Department of Agriculture, Animal and Plant Health Inspection Service, Wildlife Services, National Wildlife Research Center, 4101 LaPorte Ave., Fort Collins, Colorado, 80521, USA

## Abstract

**Context:** Ethyl-Iophenoxic acid (Et-IPA) has been widely used as a bait biomarker to determine oral bait consumption by vertebrate wildlife species. Oral bait vaccines have been delivered to numerous wildlife species to protect them from disease. The Tasmanian devil (*Sarcophilis harrisii*), the largest extant carnivorous marsupial species, is threatened by the transmissible cancers known as devil facial tumour disease (DFTD). Development of a protective DFTD vaccine is underway, and an oral bait has been proposed to deliver the vaccine in the wild. The bait delivery system requires a biomarker that can be detected for several months post-consumption in Tasmanian devils.

**Aim:** To determine the suitability of Et-IPA as a bait biomarker in the Tasmanian devil.

**Method:** Two Tasmanian devils were fed 50 mg Et-IPA (4.5 to 7.1 mg Et-IPA/kg bodyweight). Liquid chromatography with tandem mass spectrometry (LC-MS/-MS) was used to directly measure Et-IPA in baseline serum samples and samples collected on days 1, 14, 26 and 56 post-baiting.

**Key results:** Both devils retained serum Et-IPA concentrations at two orders of magnitude above negative control sera when this study concluded.

**Conclusions:** Et-IPA is a useful bait biomarker for Tasmanian devils and can be included in future DFTD bait vaccine field trials to determine bait vaccine uptake.

## INTRODUCTION

Vaccines, medications and toxicants have been delivered to vertebrate wildlife in baits for at least 50 years to prevent or treat disease, or to suppress populations of invasive or unwanted species (Baer 1976, Qureshi *et al*. 1994, Philip 2020). Central to understanding the efficacy of these control methods is the ability to monitor bait consumption. A variety of biomarkers, in particular tetracycline, rhodamine B, and derivatives of iophenoxic acid have been included in baits to identify bait consumption by target and non-target species.

Biomarkers should be non-toxic, palatable, easily incorporated into a bait, and detectable by a minimally invasive method and for a useful length of time post-ingestion (Jacoblinnert *et al*. 2021). Tetracycline and tetracycline-derivatives such as doxycycline are antibiotics that accumulate in the teeth and bones of animals when ingested and have been identified for up to 150 days post-ingestion (Van Brackle *et al*. 1994). Their use as biomarkers relies on invasive methods i.e. tooth extraction or culling, for detection. Release of antibiotics into the environment is another drawback associated with tetracyclines’ use. Rhodamine B has been widely used as a bait biomarker in mammals and can be detected in whiskers and hair for between 28 and 196 days in different species (Fisher 1999). Limitations associated with rhodamine B include palatability and difficulty of use in large production facilities (Linton Staples pers. comm.).

Ethyl-Iophenoxic acid (Et-IPA), a widely available derivative of IPA, has a long history as a useful bait biomarker in a wide variety of mammalian species. Serum Et-IPA is detected indirectly by measuring levels of protein bound iodine, or directly using liquid chromatography-mass spectrometry (LC-MS), or liquid chromatography with tandem mass spectrometry (LC-MS/-MS). These methods have detected Et-IPA in different species for up to 273 days (Ballesteros *et al*. 2013). Et-IPA has previously been tested in only two marsupial species, swamp wallabies (*Wallabia bicolor*) (Fisher *et al*. 1997) and brushtail possums (*Trichosurus vulpecula*) (Eason *et al*. 1994). The separate studies used the indirect method of Et-IPA detection, and both found plasma iodine levels had returned to baseline levels by day 7. The authors concluded that marsupials might have a physiological pathway that differs to eutherian mammals and results in faster excretion of Et-IPA (Fisher *et al*. 1997). Both studies concluded that et-IPA was not a useful biomarker in marsupials because of the short window of detection. Wallabies and possums are from the order *Diprotodontia*, which are largely herbivorous. No Et-IPA trials have been conducted to date on the carnivorous marsupial order *Dasyuromorphia*.

The Tasmanian devil is a carnivorous marsupial (Figure 1) threatened by the transmissible cancers that cause devil facial tumour disease (DFTD). Research is underway to develop an oral bait platform on which to deliver a protective devil facial tumour disease vaccine (Flies *et al*. 2020). The proposed DFTD oral bait vaccine draws on many aspects of the comprehensive oral rabies bait vaccine programs that use oral bait vaccines to immunise wildlife rabies reservoirs (reviewed in (Müller *et al*. 2015). This includes the use of IPA as the bait vaccine biomarker (Berentsen *et al*. 2019).

**Figure 1.**
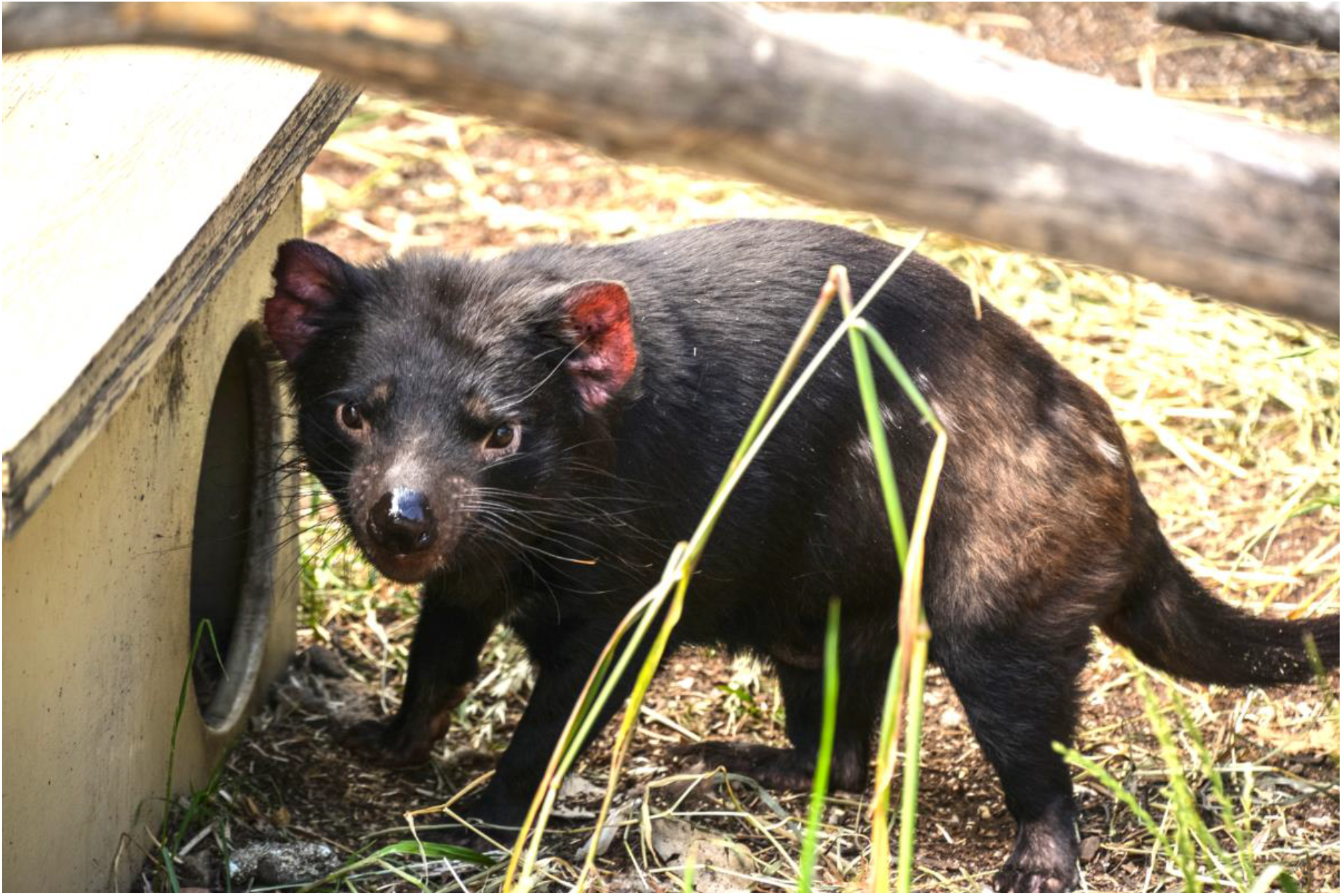
Photo of Tasmanian devil (*Sarcophilus harrisii*) (photo credit Greg Woods)

In this study we tested the suitability of Et-IPA as a bait marker in Tasmanian devils, using LC-MS/-MS to directly measure serum Et-IPA. The LC-MS/-MS method described here is similar to recently published methods in (Berentsen *et al*. 2019) and (Berentsen *et al*. 2020).

## METHODS

### Animals, IPA administration and blood sample collection

Two captive Tasmanian devils, housed individually in enclosures in accordance with the guidelines of the Department of Primary Industries, Parks, Water and Environment of Tasmania were used in this study. TD1 was a 4 year old male, body weight 11 kg, and TD2, a 5 year old female, body weight 7 kg. The devils were housed individually in 100 m^2^ enclosures. Water was provided *ad libitum* and devils fed once daily. Their diet was predominantly possum and wallaby meat, with occasional chicken and rabbit meat. All animal procedures were approved by the University of Tasmania Animal Ethics Committee under 23220 and A0017550.

200 mg Et-IPA (a-Ethyl-3-hydroxy-2,4,6-triiodohydrocinnamic acid, CAS 96-84-4, Sigma Aldrich) was dissolved in 4 ml edible corn oil, heated to 90°C to aid dissolution, then returned to room temperature. Oral baits were prepared by inoculating 1 mL of the solution (containing 50 mg Et-IPA) into a dead day-old chicken (*Gallus gallus*) and fed to each devil on day 0. The dose range of Et-IPA administered to the devils was therefore between 4.5 mg/kg and 7.1 mg/kg body weight.

The baseline blood sample was collected from each devil more than 3 months prior to the Et-IPA administration, and subsequent blood samples were collected on days 1, 14, 26 and 56 post-baiting. Blood collection was carried out under general anaesthesia following methods described in (Tovar *et al*. 2017). 3 mL blood was collected from the jugular vein into clot activating tubes. After a minimum of 30 minutes to allow for clot formation, samples were centrifuged at 1700 G for 5 minutes and the serum pipetted into 1.5 mL eppendorf tubes and stored at −80°C. Serum from a separate captive devil, housed under conditions described above, had previously been collected and stored at −80°C, and was used for the external calibration step of the Et-IPA serum detection method. Serum samples were analysed as they were collected rather than at the end of the trial.

### Et-IPA detection in serum

#### Sample Preparation

Serum samples were extracted based on the protocols of (Bruce *et al*. 2009). 200 µL of serum was taken and extracted with 800 uL of MeOH:Acetone (1:1, v/v). The mixture was vortexed for 30 seconds then placed at −20°C for 30 minutes to aid in protein precipitation. The solution was then centrifuged at 15,000 rpm for 12 min at 4°C, and the supernatant transferred to an ultra-performance liquid chromatography (UPLC) vial for analysis. Blank extractions were included with samples, together with a baseline sample recovery (spike) sample. The spike sample was produced from baseline serum, where 50 µL of the extraction solution was substituted with a 200 ng/mL working standard of Et-IPA.

For quantitation of Et-IPA in serum samples, a matrix matched external calibration was used. Two serum samples from a non-treatment devil were utilised (neat and high spike). The neat serum sample was extracted as described above. The high spike sample was extracted where 300 µL of the extraction solution was substituted with a 2000 ng/mL working standard of Et-IPA. Proportions of the neat and high spike serum extracts were then combined in various ratios to produce 8 matrix matched external calibration standards over the range 0.5 to 5000 ng/mL Et-IPA in serum. Where measured serum concentrations exceeded the external calibration range, sample extracts were diluted with baseline serum extract and reanalysed.

#### Sample Analysis

Sample analysis was undertaken using a Waters Acquity® H-class UPLC system (Waters Corporation, Milford, MA). Chromatography was performed using a Waters Acquity VanGuard C18 pre-column (5 × 2.1 mm) coupled to a Waters Acquity C18 column (100 × 2.1 mm × 1.8 μm particles). The UPLC was coupled to a Waters Xevo TQ triple quadrupole mass spectrometer (Waters Corporation). The UPLC was operated with a mobile phase consisting of 1.0% (v/v) Acetic acid in water (Solvent A) and Acetonitrile (Solvent B). Elution utilised a gradient program beginning with 50% B held for 0.5 min, moving to 95% B at 4.0 min, held for 1.0 min. At 5.5 min, composition was returned to 50% B and the column equilibrated for a further 3 minutes. The flow rate was 0.35 mL/minute and the column was held at 35°C. Injection volume was 2 µl. Typical retention time for Et-IPA was 3.0 min.

Analyses were undertaken using simultaneous single ion monitoring (SIM) and multiple reaction monitoring (MRM) in negative electrospray ionisation mode. Electrospray ionisation was performed with a capillary voltage of 2.8 kV, and individually optimised cone voltages and collision energies for each MRM transition, as described below. The desolvation temperature was 450°C, nebulising gas was nitrogen at 950 L/h and cone gas was nitrogen at 50 L/h. SIM analysis for IPA utilised the deprotonated molecule [M-H]- (m/z) 570.70 with a cone voltage of 15 V. MRM transitions monitored for IPA were [M-H]- (m/z) 570.70 to 442.80 (cone voltage 15 V; collision energy 17 V) and (m/z) 570.70 to 126.80 (cone voltage 15 V; collision energy 13 V. Dwell time per channel was 161 ms.

## RESULTS

Both of the devils readily accepted and consumed the Et-IPA marked baits. TD1 consumed the entire piece of Et-IPA laden meat within 30 minutes of it being offered and TD2 consumed it immediately.

External, matrix matched calibration was successfully demonstrated over the concentration range 0.5 to 5000 ng/mL of Et-IPA in devil serum. A linear correlation coefficient of 0.9974 +/- 0.0013 (n=3) was achieved, with an example shown in Figure 2. Recovery of Et-IPA from a 50 ng/ml spike into blank devil serum was achieved at 70 +/- 3% (n=3) over the course of three independent (inter-day) analyses.

**Figure 2.**
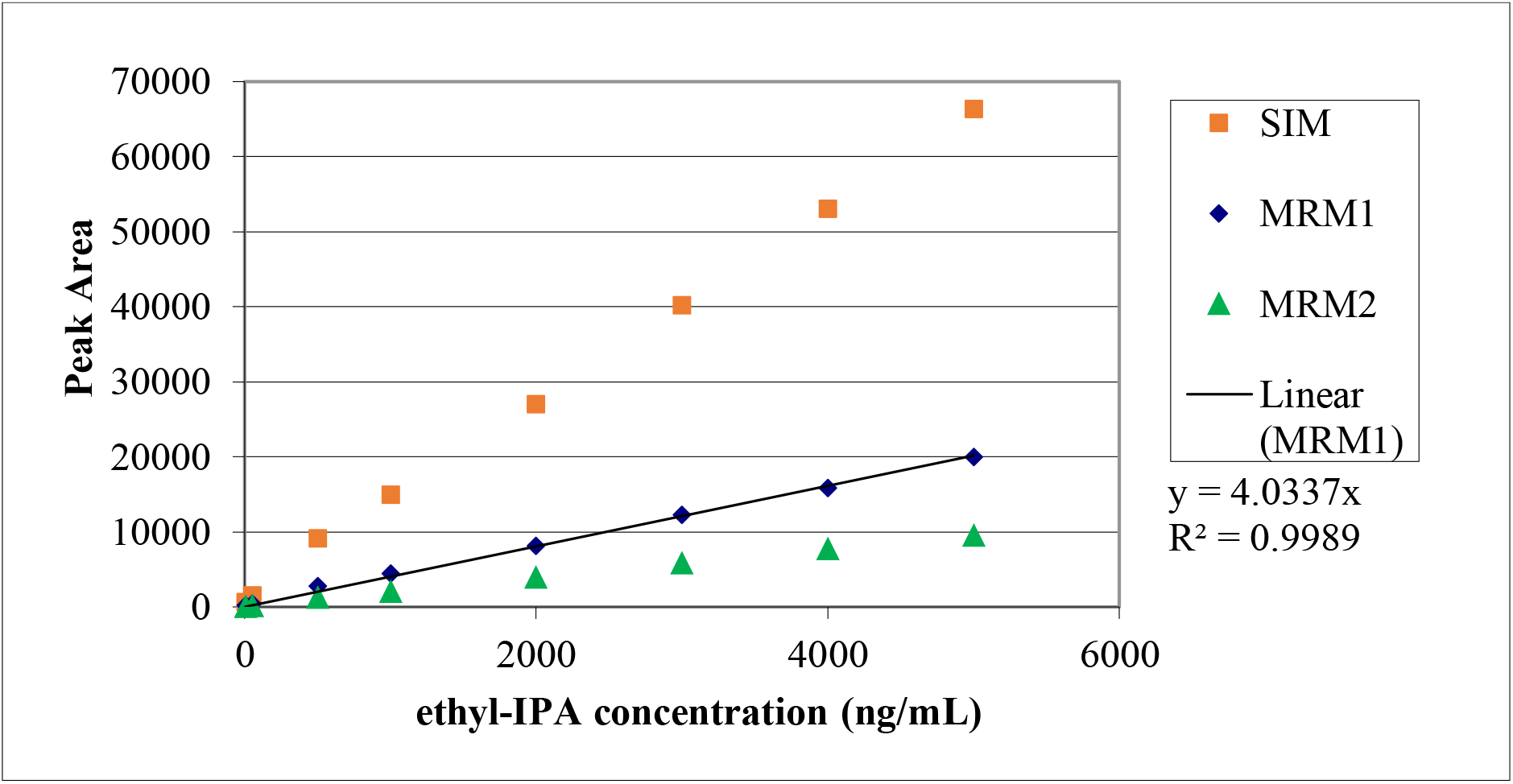
Three modes of analysis for detecting Ethyl-IPA in Tasmanian devil serum samples (day 1). First, single ion monitoring (SIM) detection of the Et-IPA precursor deprotonated molecular ion [M-H]- (m/z) 570.7. Second and third are the multiple reaction monitoring (MRM) 1 & 2 molecular transitions, where the precursor ion is then fragmented into two specific product ions, (m/z) 442.8 and 126.8 respectively. The MRM1 data is that used to undertake the actual calibration, and this is the data that is displayed for the samples.

Analysis of baseline serum samples for both TD1 and TD2 demonstrated no sample analysis interferences (Figure 3A). The highest concentration of serum Et-IPA (13,624 ng/mL) was observed for TD1 on day 1, post-ingestion (Figure 3B). This was high enough to require dilution of the serum extract with baseline serum extract, to maintain the appropriate matrix effect and get the sample into the calibration range.

**Figure 3.**
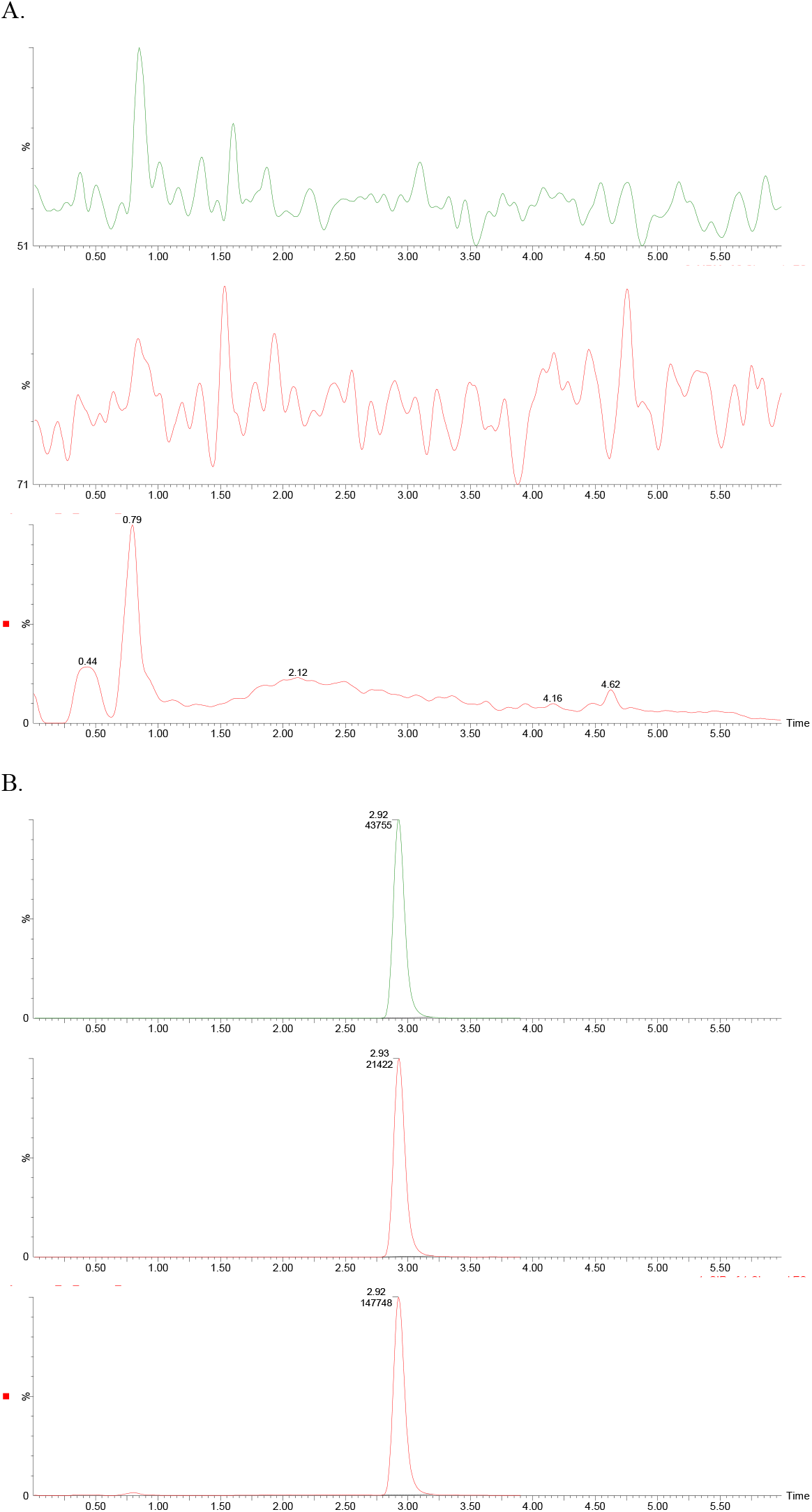
Chromatograms of serum from devil TD1: baseline (panel A), and 1 day post-ingestion of Et-IPA (panel B). For both panels, upper chromatogram represents MRM1 [M-H]- (m/z) 570.7 to 442.8. Middle chromatogram represents MRM2 [M-H]- (m/z) 570.7 to 126.8. Lower chromatogram represents SIM (m/z) 570.7.

Both devils retained quantifiable levels of serum Et-IPA at day 56 post-baiting when this study concluded (TD1 735 ng/mL; TD2 94 ng/mL, Figure 4).

**Figure 4.**
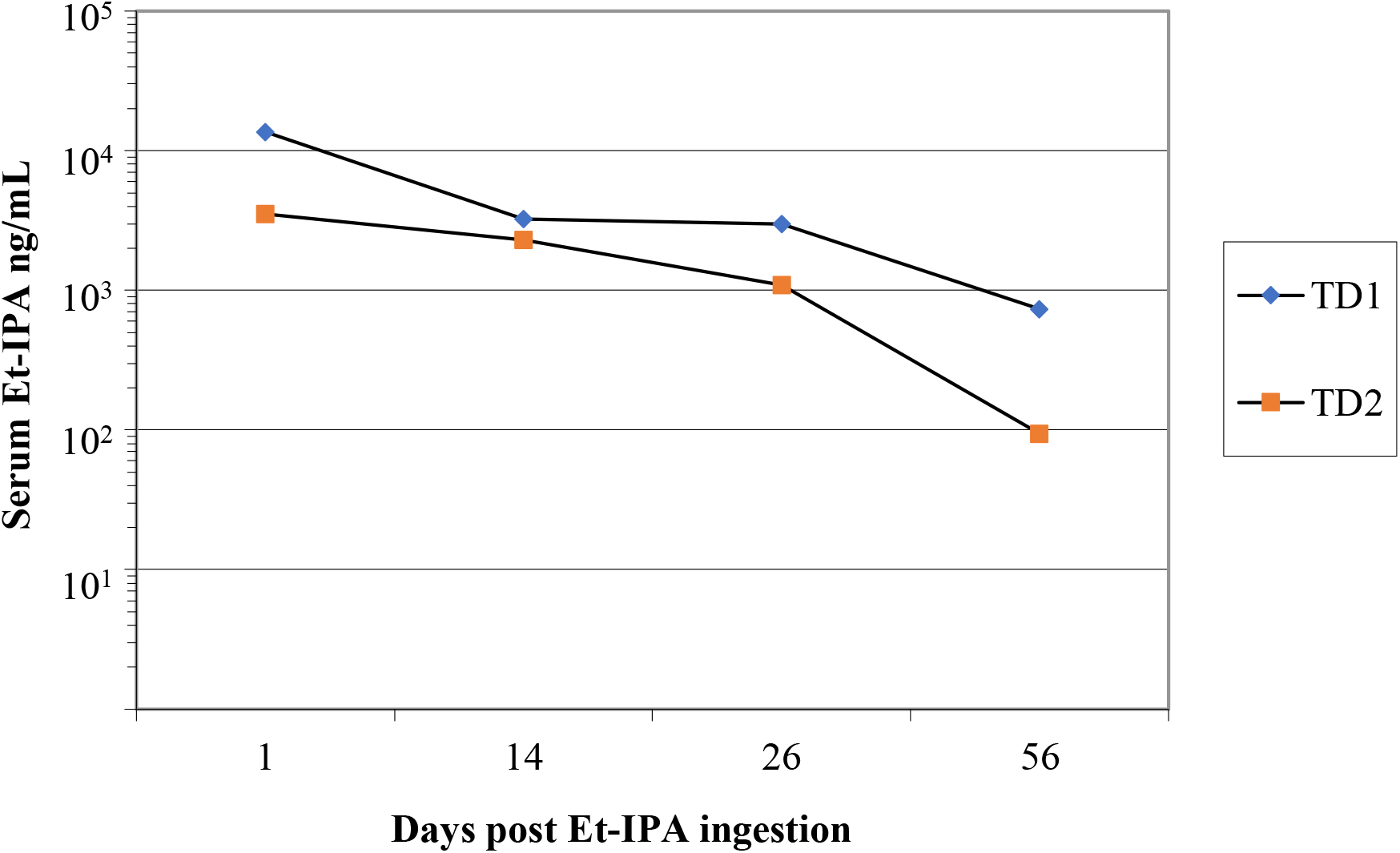
Serum concentration of Et-IPA in TD1 and TD2 over 8 weeks post-ingestion of Et-IPA. Serum concentrations were measured in ng/mL and shown on log_10_ scale

## DISCUSSION

This study is the first to demonstrate that Et-IPA is a useful biomarker in a marsupial species. We used the LC-MS/-MS method to directly detect Et-IPA in the serum of Tasmanian devils for eight weeks following oral administration of between 4.5 and 7.1 mg/kg Et-IPA. In previous studies, serum iodine was not detectable in two other marsupial species, swamp wallabies and brushtail possums, 7 days after Et-IPA ingestion despite administration of relatively high doses of Et-IPA (up to 30 mg/kg for wallabies). It was hypothesized the short window of iodine detection was due to a difference in Et-IPA metabolism between marsupial and eutherian mammals (Fisher *et al*. 1997). Those studies used the indirect method of serum iodine measurement to determine the duration of Et-IPA as a biomarker in contrast to the more sensitive LC-MS/-MS method described here.

Tasmanian devils are carnivorous in contrast to the herbivorous wallabies and omnivorous possums, which may have some effect on Et-IPA metabolism. We suggest that the LC-MS/-MS method may also detect Et-IPA in these species for a useful length of time. We expect that Et-IPA would be detected in devils beyond 56 days (when this study concluded) due to the serum Et-IPA in the twos devils at day 56 measuring two orders of magnitude above the minimum in the standard curve (0.5 ng/mL). Furthermore, the serum Et-IPA concentrations in each devil decreased by approximately 1.5 orders of magnitude over the 2-month study.

The LC-MS/-MS methods described here used a similar instrumental approach to that described previously in (Berentsen *et al*. 2019), but we used a different experimental protocol, most notably with our approach to quantitation. The (Berentsen *et al*. 2019) study, which detected and differentiated between Et-IPA and methyl-IPA in small Indian mongoose serum samples, used external calibration in a water matrix and then corrected for recovery and matrix effect by the use of methyl-IPA as the internal standard. Our approach was to use fully matrix matched external calibration of Et-IPA in devil serum. This showed no Et-IPA in the negative controls but a clear signal in serum containing the Et-IPA spike and serum from devils consuming Et-IPA.

Methyl-IPA, another derivative of IPA that differs to the more widely used Et-IPA has been incorporated into field trials that require distinction between the timing and/ or location of bait uptake (Berentsen *et al*. 2020). We could not identify any commercial suppliers of methyl-IPA at this time, precluding its broader use for temporal and spatial bait uptake field studies. Rhodamine B has been trialled in Tasmanian devils (Bell *et al*. 2020) and was detectable in whiskers when tested a minimum of two weeks post-ingestion. If the limitations of rhodamine B i.e. palatability and ease of incorporation into bait matrix, can be overcome, it could be used in combination with Et-IPA for such studies. Future oral DFTD bait vaccination trials for Tasmanian devils that include an immune priming component followed by an immune boosting component would benefit from two different biomarkers.

## Acknowledgements

We thank Ginny Ralph for taking care of the captive devils used in this trial.

